# A diverse gut virome from *Drosophila melanogaster*

**DOI:** 10.1101/2024.03.19.585549

**Authors:** Mina Hojat Ansari, Fabian Staubach, Nurper Alacatli, Darren J Obbard

**Affiliations:** Department of Evolution and Ecology, University of Freiburg, Freiburg, Germany; Institute of Ecology and Evolution, University of Edinburgh, Edinburgh EH9 3FL, United Kingdom

**Keywords:** gut microbiome, DNA virus, phage, Drosophila

## Abstract

*Drosophila melanogaster* is not only one of the most important models of antiviral immunity in invertebrates, but is also a powerful model for research of the gut microbiome. Although recent studies have continued to improve our knowledge of the fly gut microbiota, the viral component of the microbiome has remained unexplored. Here we explore the viral component of the *Drosophila melanogaster* gut microbiome using deep metagenomic DNA sequencing. We recovered 3035 phage sequences, resulting in 167 viral Metagenome-Assembled Genomes. The majority of these sequences are potentially novel bacteriophages from the order *Caudovirales*, which mainly target major gut bacteria of *D. melanogaster*, including *Lactobacillus*, *Acetobacter*, and *Gluconobacter*. Our functional annotation and discovery of auxiliary metabolic genes showed that these bacteriophages have the potential to influence microbial metabolism and genetic information processing. We also identified evidence of known fly pathogens Drosophila Kallithea nudivirus, Vesanto bidna-like virus, and Viltain densovirus, some of which were common in our studied populations. Our findings reveal a complex and diverse phage community in the *D. melanogaster* gut microbiome, paving the way to study host-phage related research in the natural microbial communities.

## Introduction

The study of viruses, including bacteriophage (‘phages’; bacteria and archaea-infecting viruses) has recently garnered increasing interest in scientific communities due to the advancement of metagenomic techniques that facilitate the identification of viral genomes. Bacteriophages have been founded in nearly all investigated ecosystems and have been recognized as the most abundant and diverse biological entities on Earth [1]. They play an important role in microbial communities by affecting microbial diversity and metabolism [2, 3], and mediating phage-born antibiotic resistance within hosts [4].

Studies have also revealed the presence of auxiliary metabolic genes (AMGs) in bacteriophages [5]. These genes encode proteins that can regulate metabolic processes within host cells. In addition, bacteriophages have been found to regulate transcription and translation [6], and influence the decision to lyse or enter a lysogenic state [7], further underscoring the importance of characterizing bacteriophages and their interactions with their hosts.

In humans, it has been demonstrated that bacteriophage are mechanistically involved in human health and disease [8]. However, information about phages as an important component of microbiome infecting most species, is still lacking. This is also true for the well-studied model organism *Drosophila*, despite being a powerful model for gut microbiome research [9, 10] and the study of evolution and ecology of host-microbe interactions [11, 12].

The fruit fly *D. melanogaster* is a model organism that naturally interacts with a variety of microorganisms due to its feeding, mating, and ovipositing on fermenting and rotten fruits. *Drosophila melanogaster* has not only been established as a powerful model for gut microbiome research [9, 10], but is also well known as an important model for invertebrate antiviral immunity [13, 14]. However, most studies aimed at describing viruses in *Drosophila* have focused on the detection of RNA viruses of the host [15, 16], and studies on the diversity of DNA viruses have exclusively described eukaryotic DNA viruses [17]. Thus, almost nothing is known about the diversity or function of prokaryotic viruses—bacteriophages— associated with *Drosophila*. Here we aim to provide an initial characterization of the prokaryotic viral microbiome associated with *D. melanogaster* by applying metagenomic sequencing to screen the guts of wild-caught flies for viruses.

## Methods

### Sample Collection and Metagenomic Sequencing

A total of 851 male *Drosophila melanogaster* were collected from four sampling locations in Germany (Baden-Württemberg: for more details see Supplementary Fig. S1), over a two-month period between late August and early October-2019. For sample collection, we used traps baited with a mixture of apple and cherry, following a standardized protocol [18]. The flies were anesthetized on a fly pad using CO_2_ and subsequently transferred into empty Eppendorf tubes. The Eppendorf tubes were then dipped into liquid nitrogen to freeze the samples. Following this, all fly samples were dissected to extract the gut, and then the individual gut samples were stored in PBS at -80°C until DNA extraction.

We extracted DNA from each gut sample individually following the DNA extraction protocol described in detail in [18] with slight modification. Briefly, the samples were homogenized using a bead beater (Qiagen Tissue Lyzer II), with the addition of lysozyme to improve the efficiency of nucleic acid extraction. Protein was digested with Proteinase K, and RNA was removed using RNAse. We precipitated DNA using a combination of Phenol-Chloroform-Isoamyl alcohol and after washing with alcohol the DNA was resuspended in nuclease-free water. The NEBNext® Multiplex Oligos with unique dual index primer pairs for Illumina was used for library preparation. Then samples were sequenced on the Novaseq 6000 platform with 150bp paired end sequencing strategy at the Life & Brain research centre (University Hospital Bonn).

### Metagenome assembly

We used Cutadapt [19] and FastQC [20] implemented in TrimGalore v0.6.6 [19] to trim adapters and control the quality of the raw reads. Then, to remove human and host sequences, we mapped the genomic DNA of the data from each of the 851 sample to the host *D. melanogaster* and the human genome (hg38), using bbmap [21]. Finally, we assembled the remaining paired-end reads using MEGAHIT [22]. To improve the assembly of low-abundance components of the microbiome, we co-assembled reads from multiple individuals, pooling by sampling location, for a total of four co-assemblies.

### Recovering high-quality viruses and Binning of viral genomes

We predicted viral contigs among metagenome contigs using VIBRANT v1.2.1 [23] with the default parameters. VIBRANT uses machine learning and a protein similarity approach to accurately recover bacteriophages [24]. In outline, VIBRANT uses the contigs predicted by Prodigal v2.6.3 [25] and annotates them with the Kyoto Encyclopedia of Genes and Genomes (KEGG) KoFam [26], Pfam [27], and virus orthologous group (VOG) [28] databases using HMMER v3.3.2 [29]. At the same time, we also inferred auxiliary metabolic genes (AMG) using VIBRANT. VIBRANT employs neural networks of protein signatures, and a novel v-score metric to overcome conventional limitations, enabling comprehensive identification of lytic viral genomes, thereby capable of providing estimates of genome quality and differentiation between lytic and lysogenic viruses.

The completeness and quality of all phage contigs were assessed using CheckV v1.4 [30] with the parameters “end_to_end” and removing all contigs shorter than 3kb. To ensure the quality of retained phage contigs, we ran VirSorter v2.2.3 [31] and DeepVirfinder (v. 1.0) [32] against the same assembled data. We considered the phage contigs recovered by VIBRANT as true bacteriophages if the prediction scores of VirSorter2 (groups include-dsDNAphage, ssDNA viruses, NCLDV-Nucleocytoviricota) and DeepVirfinder also were above ≥ 0.5.

DeepVirFinder has especially been advocated for the identification of short (1< kbp) phage genome fragments [33]. The dsDNA, ssDNA and NCLDV, respectively, represent double-strand and single-strand DNA viruses and nucleo-cytoplasmic large DNA viruses. However, to reduce the chance of false positives given the sensitive cut-off point for the length of the VIBRANT and VirSorter2 fragment of 3 kbp, the short contigs <3 kbp) were also retained if they had been recovered by all three pieces of software and had VirSorter2 and DeepVirfinder quality scores ≥ 0.9. Contigs with provirus contamination were also retained if they exhibited a clear shift from the host genome and met the aforementioned criteria, after extraction of the proviral region.

All identified contigs from the four co-assemblies were pooled and dereplicated (–method longest) using vRhyme v.1.1.0 [34]. The quality of the final nonredundant phage sequences has been assessed using CheckV v1.4 [30]. We also used Cenote-Taker 2 v2.1.5 with “-am True” [8] to further assess and annotate the nonredundant recovered sequences.

Finally, we used the nonredundant viral contigs binning strategy to group scaffolds into putative genomes using vRhyme [34] with the default parameters. As part of bin construction, vRhyme checks for protein redundancy and this metric can be used for post-binning filtration (Redundant protein <1 unlikely to be contaminated: 2<5 may not be contaminated; >6 are more likely to be contaminated) [34].

To perform a quantitative analysis of the obtained phage contigs, we used metaWRAP with “quant_bins” module to map the high-quality reads against the recovered contigs. Then the average abundance of each (viral metagenome-assembled genome) vMAG in each of 851 sample was calculated using Salmon [35], reporting quantities as transcripts per million (TPM) [36] and taking the length-weighted average of the contig abundances.

We performed principal coordinate analysis (PCoA) and visualized the Bray–Curtis distances of relative viral abundances at each sampling location using the vegan package (v2.5-7) in R v4.2.0 [37]. Then the observed variation was further tested using Adonis2, also Betadisper test in the vegan package [38], to gain a better understanding of the underlying patterns and structure in the data set as well as validate the findings obtained through PCoA. Adonis2 helps determine if the observed groupings in PCoA are statistically significant, while Betadisper provides insights into the within-group variability and helps identify if the groups identified by PCoA have significantly different spread or variability. We also made a heatmap to visualize the differences between phage composition of different localities using Seaborn clustermap v 0.12.2 [39].

### Protein clustering and taxonomy of viral contigs

We evaluated viral contig similarity and constructed a viral protein clustering network with vConTACT2 0.11.3 [40] using the viral Refseq v211 database. The vConTACT2 method uses gene-sharing networks to assign the taxonomy of viruses based on their sequence. The viral sequences are then classified as ‘clustered’ (high-confidence clustering), ‘overlap’ (sharing overlap gene content), ‘outlier ‘(weakly associated with a cluster) or ‘singleton’ (few or no shared gene content) [40]. The network was visualised using Cytoscape v3.9.1 [41]. We performed host assignment and taxonomic annotation at the family and genus level with VPF-Class [42] using VPF classified files from 17 August 2022 to build the reference database. For preliminary taxonomic annotation, we considered a VPF membership ratio cutoff of 0.2 and 0.3 at the family and genus level, respectively.

### Host Assignment of recovered bacteriophage contigs

To assign viral genomic sequences to their specific host, we applied two strategies. First, we used VPF-class tool, which takes advantage of the Viral Protein Families (VPFs) and creates a protein-based database from the IMG/VR database. The VPF-class tools then conducts a protein family search for each query genome and concludes the host prediction based on the distribution of these VPFs in reference phage genomes, first at the domain and then at the host family and genus levels [42]. In the second approach, we used CRISPR spacer-based bacterial host predictions strategy [43]. The CRISPR spacer-based approach uses information from CRISPR–Cas, which works as adaptive immune systems, and has access to a database of more than 11 million spacers [43]. We therefore searched the spacer database with our viral sequences using blastn and predicted bacterial hosts at the genus level on the basis of sequence similarity.

### DNA viruses of Drosophila melanogaster

Initial taxonomic assignment of the recovered contigs suggested that at least one nudivirus was present. We therefore used blastn to test whether this was a previously known viral pathogen of *Drosophila*. To detect the presence of other previously-known DNA viruses of Drosophila, we mapped the high-quality short reads to the reference genomes of Drosophila Kallithea nudivirus, Drosophila Viltain densovirus, Drosophila Linvill Road densovirus, and Drosophila Vesanto bidna-like virus to quantify the amount of each virus in each fly [15, 17, 18]. We set a threshold of 1 per cent of the fly genome copy number to define infection status, and using these data, we were able to estimate the prevalence of these fly pathogens in our 851 samples.

## Results

### The metagenomic virome of *D. melanogaster* gut

We processed 851 metagenomes generated from *D. melanogaster* gut samples, resulting in nearly 38 billion high quality reads (range 1 to 90 million per sample). After mapping to the host, we retained 7.7 billion non-fly reads, and from these VIBRANT recovered 9124 metagenomic viral contigs (MVCs) that ranged in length from 1 to 545 kb with a mean length of approximately 7kb, and a total of 4964 contigs greater than 3Kb (Supplementary Table S1). After comparing the result with the finding from two other pieces of software (VirSorter2 and DeepVirfinder), filtering and dereplication, we identified 3040 unique contigs (Supplementary Table S2). Of the 3040 non-redundant contigs, 940 contigs were shorter than 3kb, and 1145 contigs were greater than 5kb. VIBRANT annotation identified 30 out of the 3040 nonredundant viral as complete circular genomes, 31 with high quality, 79 with medium-quality and 2900 as low-quality draft. Cenote-Taker 2 was also used, which checks for direct terminal repeats (DTRs) and inverted terminal repeats (ITRs), representing two common end features of complete viral genome. The Cenote-Taker 2 showed that 32 and 16 contigs out of 3040 contained DTRs and ITRs, respectively. These data showed 30 of 32 DTR-containing contigs matched with contigs identified as complete circular contigs by VIBRANT (Supplementary Table S2).

To further evaluate the final viral sequences, we applied checkV and accordingly most of the sequences (2821) were estimated to be low-quality genome fragments (<50% complete), with only 67 with medium-quality (50–90% complete) and a total of 37 contigs were estimated to be high-quality (>90% complete), of which 20 were predicted to be completed genome, also 115 contigs were not-determined (Supplementary Fig. S2). The chance of not-determined contigs to be false positive is low, as we retained them as viruses if they have been predicted with all three pieces of software (with VirSorter2 and DeepVirfinder quality scores ≥ 0.9).

Therefore, the not-determined contigs are more likely to be too short to categorise, or the sequences are too divergent from the CheckV references—although Cenote-Taker 2 also could not predict halmarks for most of these 115 contigs. Next, we used VIBRANT to analyze whether the 3040 identified contigs were likely to have a lytic lifecycle. We found that 96.7 % of them (2941 viral contigs) were predicted to have a lytic lifestyle, while only 3.3% represented a temperate lifestyle (Supplementary Table S2).

### Protein Classification and Taxonomy of the virus contigs from the *D. melanogaster* gut

We evaluated the degree to which the newly recovered *D. melanogaster* gut-associated viruses were represented in available public databases (Fig. 1). Using a method that applies gene-sharing networks to classify sequences (vConTACT2), we determined that of 3040 non-redundant contigs over 1kb in length, 716 were singletons, 741 were outliers, 294 had genomes sharing overlap with multiple established clusters (i.e. cannot be unambiguously clustered), and 1289 were clustered (either with Refseq sequences or only with each other) (Supplementary Table S3 and Fig. S3A).

**Fig. 1:**
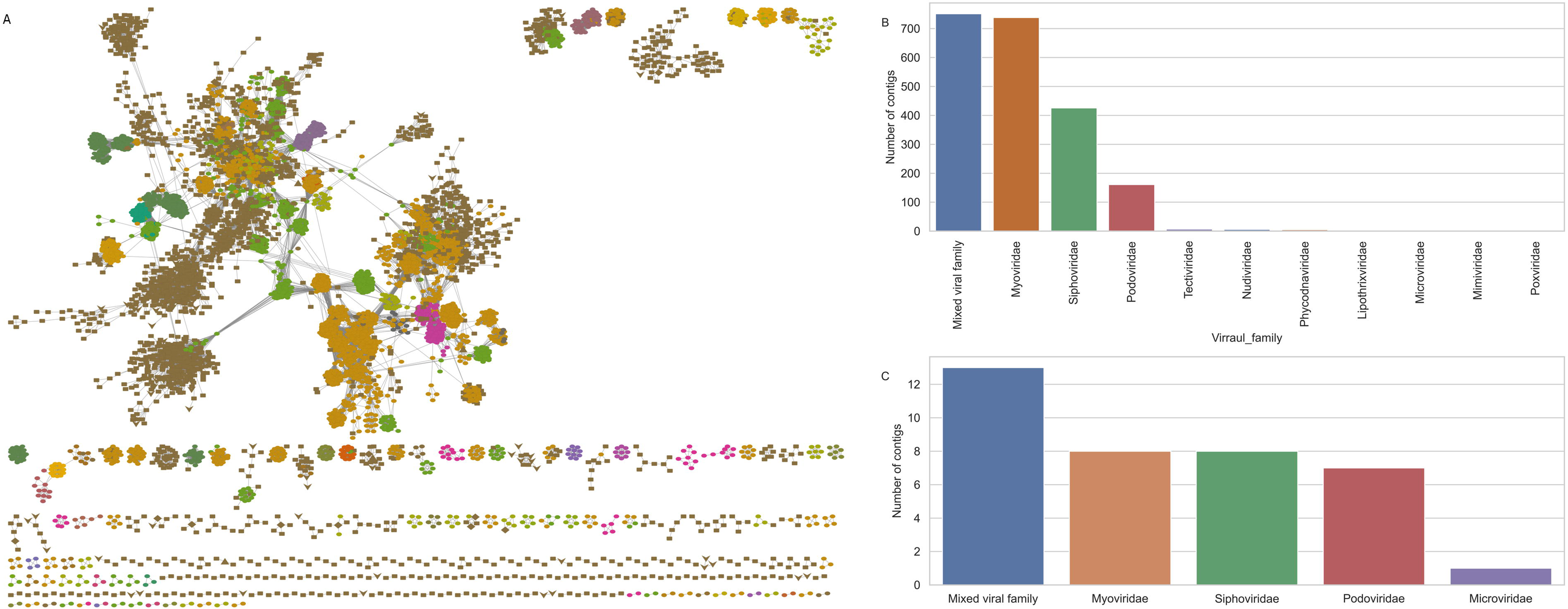
Taxonomy assignment and protein network clustering of viral sequences. A: Protein clustering network of the recovered viral sequences from *D. melanogaster* (dark-brown) compared to previously described viruses from Refseq (other colors). The most common viral families are highlighted with orange (Siphoviridae), green (Myoviridae), and olive (Podoviridae) colors. B: Taxonomy of viral sequences at family level, determined by VPF class. C: Viral families associated with high-quality viral sequences.

The 1289 clustered contigs formed 469 clusters, of which 434 clusters did not include any sequences from the Refseq database (Fig. 1A and Supplementary Table S3). The remaining 35 clusters, representing 67 contigs, were classified mainly as *Siphoviridae*, *Myoviridae*, and *Podoviridae* (Supplementary Fig. S3B). Furthermore, members of *Schitoviridae*, *Herelleviridae*, and *Autographiviridae* were present, although they comprised very small numbers of contigs (Fig. 1A and Supplementary Fig. S3B). In general, the resulting protein similarity network represented the newly identified phage sequences as broadly dispersed across the known viruses from the RefSeq database (Fig. 1A), despite the fact that 25% of the recovered sequences were also singletons (i.e. not presented in the network graph). The protein similarity network classification, however, showed that the higher-quality contigs (>50 % complete) tended to cluster more closely with previously identified viruses in the Refseq database, where almost 30% of these sequences assigned to known phages (Supplementary Fig. S4). Assignment success depended partly on contig length (Supplementary Fig. S5), with 53 (4.6%) of larger (>5kb) contigs being assigned, but only 14 (∼1%) of the smaller contigs (Supplementary Figs. S5A and S5B). However, the assignment of the large contigs to different phage families was comparable to the entire data set (Fig. 1 and Supplementary Figs. S3 versus Figs. S5).

To determine the family level taxonomy of the recovered contigs, we also characterised overlap in the composition of their viral protein families (VPF) with known viral taxa (Supplementary Table S4). Most of the recovered contigs have VPF compositions with family-level relatedness to *Siphoviridae*, *Myoviridae*, *Podoviridae,* and in lower ratios with *Tectiviridae* and *Microviridae* (an ssDNA virus). We also identified VPFs from archaeal virus families (*Lipothrixviridae*, *Sphaerolipoviridae*, *Rudiviridae*, and *Turriviridae*), NCLDV, and *Nudiviridae*—a family of DNA viruses known to infect eukaryotes, including *Drosophila* (Fig. 1B and Supplementary Table S4).

Of the 37 high-quality genomes, the VPF membership ratios of almost all suggest that they were related to bacteriophage families including *Myoviridae*, *Siphoviridae*, *Podoviridae*, and *Microviridae* (Fig. 1C). Although 27 (73%) of these almost complete phage genomes did not cluster with known reference phages (Supplementary Fig. S4), their VPF membership ratios were primarily related to *Caudovirales* (Fig. 1B and Supplementary Table S4), which may point to novel families or genera within this order.

### Host Assignment of bacteriophage contigs

To predict the hosts of the recovered phage sequences, we took advantage of the VPFs as well as CRISPR-spacer sequences. Using the VPF approaches with a membership ratio and a confidence score cutoff of 0.6 and 0.8 respectively, 650 contigs could be linked to a host domain, of which almost all were assigned to the bacterial host (Supplementary Table S5).

Further investigation showed that 267 of these phage contigs were linked to 15 host families and 154 of them could be related to 20 host genera. All identified potential hosts reflect either the gut or environmental bacteria (Supplementary Table S5). Of the 156 contigs that hit the host at genus-level, 43 could be related to the genus *Pseudomonas*, 30 to *Lactococcus*, 25 to *Staphylococcus*, 10 to *Escherichia*, and eight to *Lactobacillus* (Supplementary Table S5).

On the other hand, the CRISPR-spacer approach, which was used to predict bacterial hosts of determined phages, linked 488 bacteriophage contigs to 61 unique bacterial genera. Of these 488 contigs, most of them (>300 contigs) hit against *Drosophila* gut specific bacteria (such as 74 *Lactobacillus* hits, 66 Gluconobacter hits, 48 Acetobacter hits, 31 Acinetobacter hits, 20 Pseudomonas hits, 19 Serratia hits, 15 Corynebacterium hits, 13 Klebsiella hits, 13 Komagataeibacter hits, etc.) (Supplementary Table S5).

Comparing the result from VPF and CRISPR-spacer approaches showed 29 contigs that hit the host at genus-level shared between two approaches, but only 69% of them (20 contigs) hit the same bacteria genus using both approaches (Supplementary Table S5). Further investigation showed that the VPF tools also predicted the presence of similar genus as CRISPR-spacer for these nine contigs, but were excluded due to our inclusion threshold (including only VPFs with a membership ratio and confident score of ≥0.6 and ≥ 0.8, respectively).

### Functional Potential of *D. melanogaster* associated phage contigs from gut

Annotations from KEGG, Pfam, and VOG databases were used to gain insight into the functional potential encoded by the recovered contigs. Based on the available databases, only 50% (1549 contigs) of the contigs were assigned a function based on KEGG. However, the majority of phage associated functional genes were annotated with metabolism functions, including nucleotide, amino acid, lipid, energy and carbohydrate metabolism (Supplementary Table S6). Furthermore, functional genes associated with DNA repair, replication, and recombination, also transcription, were including quite large proportion (Supplementary Table S6).

We identified a total of 149 phage contigs that encoded at least one AMG and could be assigned to 42 unique KEGG orthologs (KO) (Supplementary Table S6). The most abundant AMGs (in terms of number of contigs involved) encoding dcm/DNMT1, folA/DHFR, mec, queE, NAMPT and folE/GCH1. These genes encode, respectively, DNA (cytosine-5)-methyltransferase 1, dihydrofolate reductase, the [CysO sulfur carrier protein] -S-L-cysteine hydrolase, the 7-cyano-7-deazaguanine synthase, nicotinamide phosphoribosyl transferase, and GTP cyclohydrolase IA (Supplementary Table 6). In other words, the AMGs derived from *D. melanogaster* tend to encode products mainly for amino acid, cofactor and vitamin metabolism and folding, sorting and degradation.

To further explore the functional potential of the identified contigs, we projected KO accession onto the KEGG pathways (Fig. 2). We used a total of 1549 phage contigs, which could be annotated based on KEGG database and assigned to 419 unique KO, for KEGG pathways projection (Supplementary Table 6). In some cases, the retrieved pathways represent functions that could directly affect bacteria, such as involvement in biofilm formation, quorum sensing, and the bacterial secretion system. In general, the retrieved pathways reflect a wide range of metabolic functions, including carbohydrate, energy, lipid, nucleotide, and amino acid metabolism. Furthermore, glycan biosynthesis and cofactors, vitamins, terpenoid and polyketide metabolism, and xenobiotic degradation were presented (Fig. 2). The identified genes in *D. melanogaster* gut phages are also involved in folding, sorting, degradation, transcription, and translation, as well as in replication and repair.

**Fig. 2:**
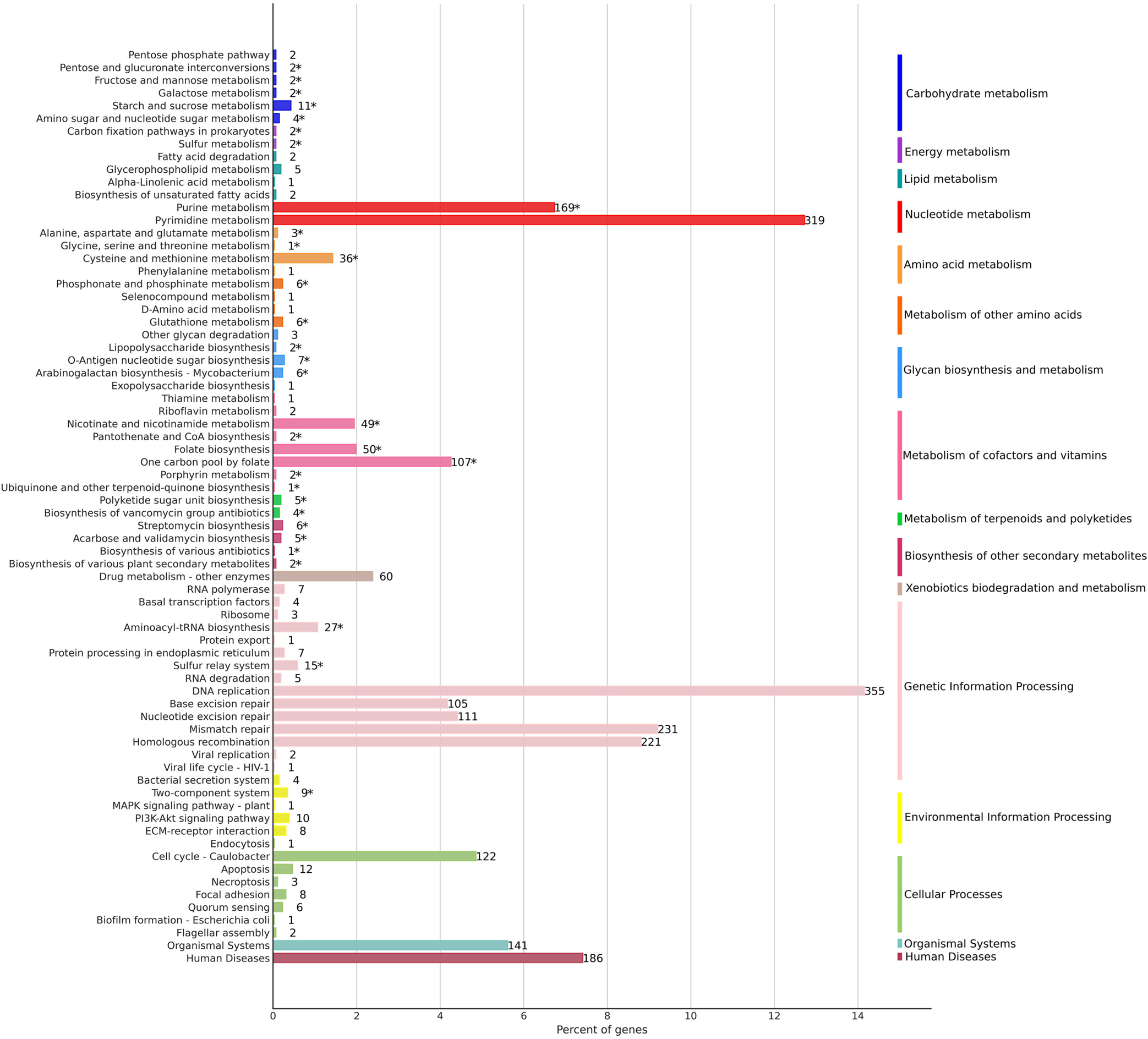
Bar plot illustrate the functional annotation results based on the KEGG pathways represented in the recovered viral sequences.

### Genome binning of phage contigs from metagenomic data from *D. melanogaster*

We used vRhyme to construct viral metagenome-assembled genomes (vMAGs), resulting in the binning of 767 contigs into 188 vMAGs (Supplementary Table 7). However, after post-binning filtration based on potential contamination of vMAGs (bins that could include sequences from multiple viral genomes), in which we filtered out bins with ≥ 6 redundant proteins, we ended up with 167 vMAGs. We then checked the relative abundance of the final vMAG sets across the collected samples and observed variation related to the sampling locations (Fig. 3A). Out of 167 vMAGs, 11 were identified in more than 70% of the samples. Among these, two genomes, which their VPF membership ratios primarily belong to the *Myoviridae* family, were detected in all 851 samples (Fig. 3B and Supplementary Fig. S6).

**Fig. 3:**
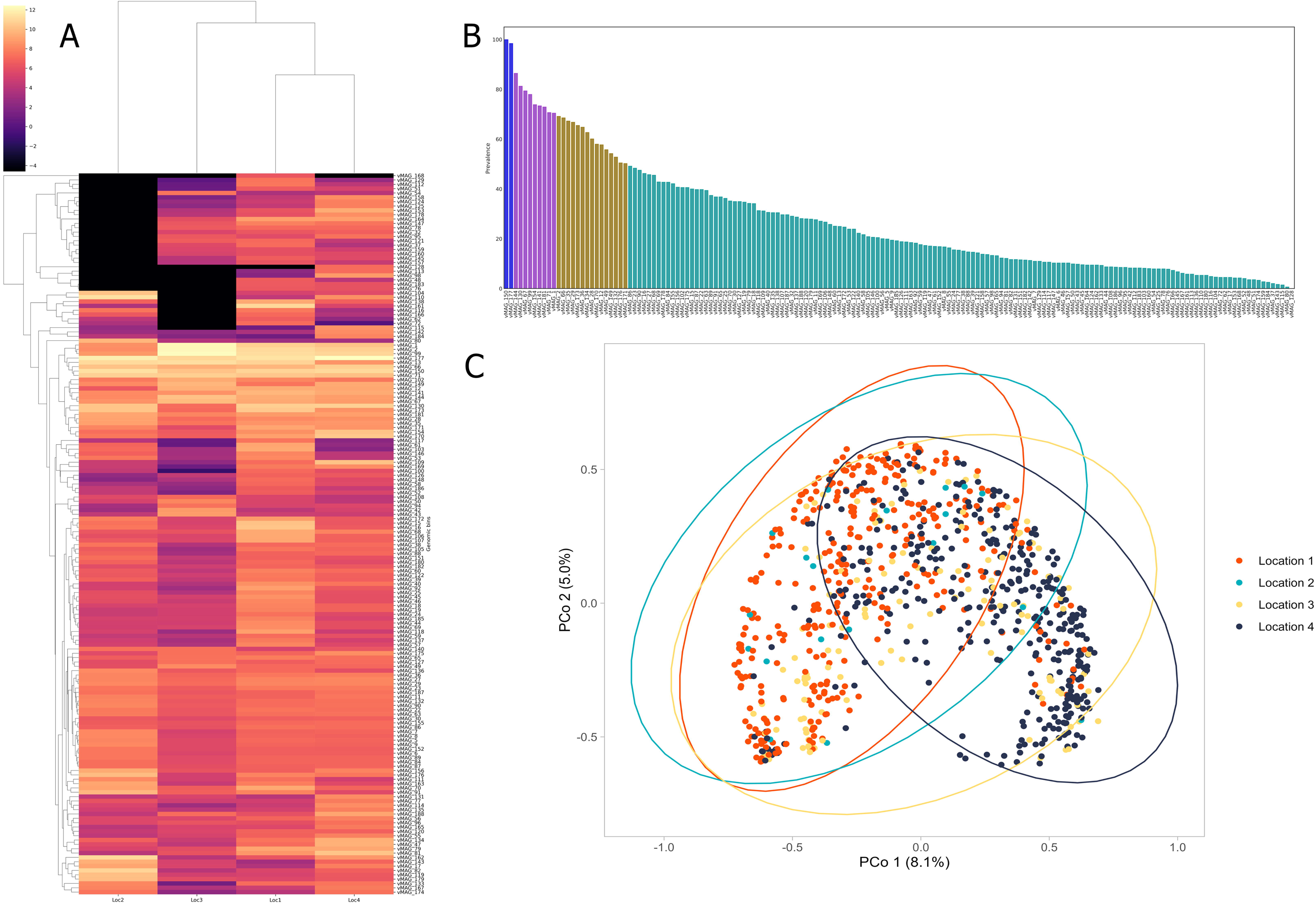
Characterization of the viral metagenome assembled genome (vMAG) recovered from *D. melanogaster* gut microbiome. A: Relative abundance and distribution of 167 recovered vMAGs from four sampling locations (Loc). B: The prevalence for each vMAG (prevalence <50%= green, 50<70%= brown, 70<98%=purple, and 99 %< showed with blue color) were illustrated in the bar chart. C: Principal component analysis using Bray-Curtis dissimilarity. The p-value and R squared values based on Adonis2 (with 10,000 permutations) were 9.999e-05, and 0.0848, respectively. The p-value for Betadisper was also significant (P= 0.001).

Next, we further investigated the pattern of viral sequences across the samples, using principal coordinate analysis (PCoA), and clustering for sampling locations was assessed using Adonis2 and Betadisper tests. A significant effect from sampling location was observed not just based on the vMAG sequences but also based on unbinned viral sequences (2381 contigs) (Fig. 3C and Supplementary Fig. S7).

### Drosophila melanogaster DNA viruses

During the analysis of virus protein composition, we identified a number of contigs deriving from nudiviruses, which are known to include multiple pathogens of *Drosophila melanogaster* (Wallace, Coffman et al., 2021). A blastn search against the NCBI viral genome database showed that these contigs were derived from Drosophila Kallithea nudivirus (NC_033829) with 99-100% coverage and a 99% identity. By mapping the high-quality short reads to the reference genome, we found that Kallithea virus occurs frequently, with a high overall prevalence of 54.2% (in 461 individuals of 851 samples with genome copy-number threshold ≥1% of the fly genome copy-number). The prevalence of Kallithea virus varied among four different sampling locations, ranging from 33.9% to 82.1%. We also detected Drosophila Esparto nudivirus, but its prevalence was extremely low, with only affected two flies.

By mapping reads to all previously-reported DNA virus genomes from *Drosophila melanogaster* [15, 17, 18], we identified two further previously-known pathogens, including Drosophila Linvill Road densovirus and Vesanto bidna-like virus. Drosophila Linvill Road densovirus was extremely rare, being detected in only seven individuals, all from sampling location 1, indicating a prevalence of 2 percent in that location and a total prevalence of 0.8% (95 % confidence intervals 0.78%-0.82%). On the other hand, *Drosophila* Vesanto virus, a multi-segmented bidna-like virus, occurred at relatively high prevalence of 20% (95% confidence interval 18-23%; 172 samples out of 851) using a threshold of ≥1% (Fig 4).

**Fig. 4:**
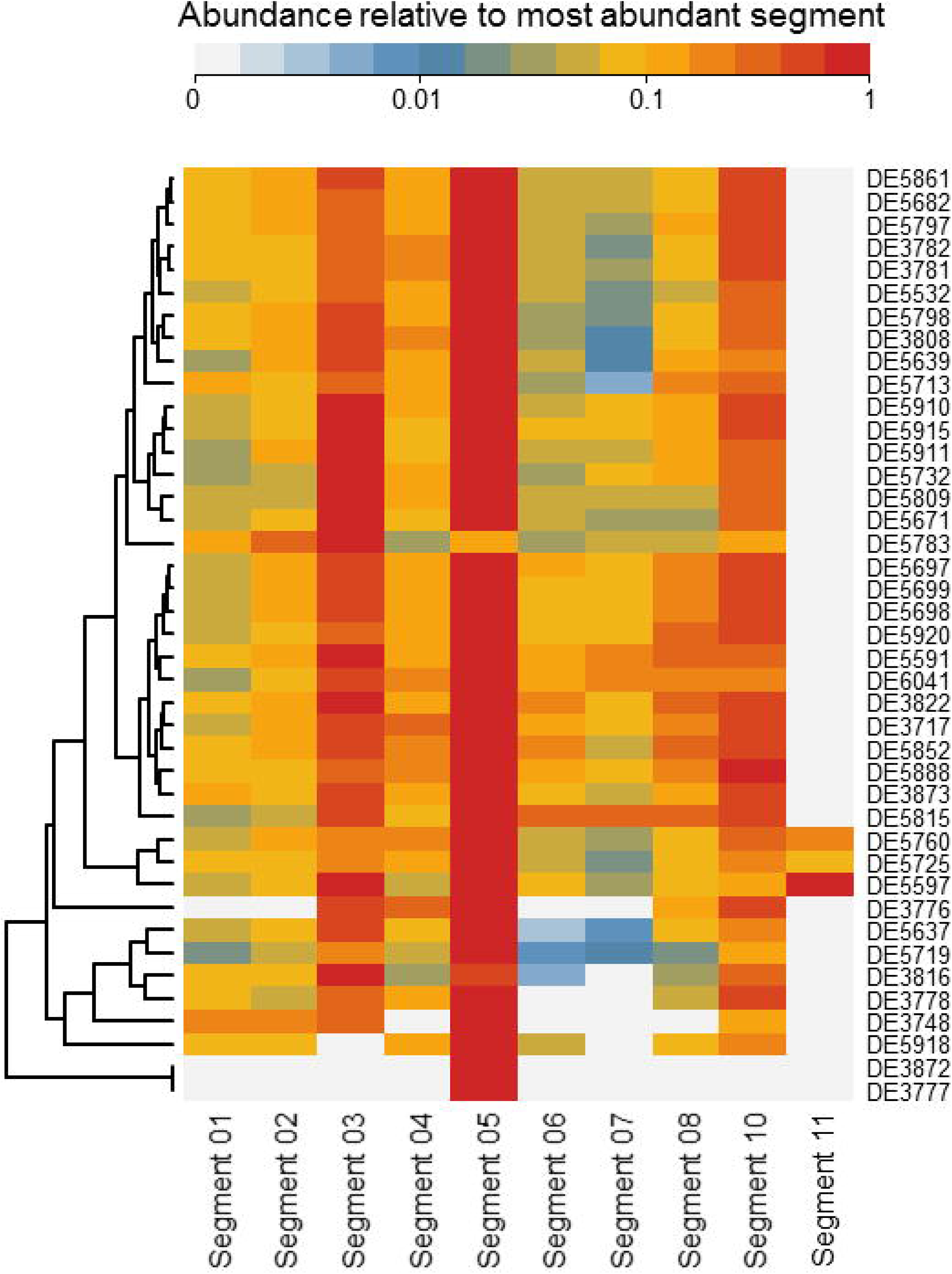
Heatmap demonestrating the relative number of sequencings reads from each of the the 10 Vesanto virus segments), for each sample with the minimum threshold of 1 percent.

Almost all previous sequencing of Vesanto virus has been based on large metagenomic pools, leaving the associations between putative segments uncertain, as many independent infections are represented in each pool [17]. Here, based on single fly sequencing, we did not detect segments 9 or 12 —which were also absent from some population pools reported in [17].

Detection of the other segments was variable, with segment 5 being detected in 165 out of 851 samples (Fig 4A, i.e. nearly all individuals infected with the Vesanto virus), segments 3 and 10 being detectable in 86 and 56 samples respectively and segments 1, 2, 4, 6, 7, and 8 each appearing in more than 35 individuals. Segment 11, was extremely rare, detected in only three samples.

Excluding segment 11, all other segments were detectable in those infected individuals that had the highest Vesanto virus read numbers (e.g., samples DE5797 and DE5682, which exhibited almost 2.5 and 1.8 million Vesanto virus reads, respectively). This is consistent with some segments (e.g. segments 9, 11, and 12) being ‘optional’ or ‘satellite’ components and other segments differing in copy-number, with segment 5 being the most readily detectable.

## Discussion

The virome is increasingly acknowledged as an essential part of the microbiome, playing a key role in coevolution with prokaryotic and eukaryotic hosts by both increasing and decreasing host fitness [44]. Here we used a metagenomic approach to elucidate the structure and function of the viral community associated with the gut of wild-caught *Drosophila melanogaster*, with a focus on prokaryotic viruses.

### Phage associated with the *Drosophila* microbiome

Our study found that *D. melanogaster* harbors a diverse community of temperate and lytic bacteriophages in their gut (Supplementary Table S2). We also observed distinct variations in the structure of *D. melanogaster* bacteriophage communities from different sampling locations. This could suggest the gut microbiome in association with the host and environment is highly dynamic [45, 46].

We used two methods for host assignment of viral sequences: the VPF identification and the CRISPR spacers similarity method. Both methods assigned a relatively low percentage of the phages to a particular host (smaller than 16% in total). This is consistent with previous findings in honeybees [47] or humans [48]. Therefore, the number of viruses that could infect members of the fly core gut bacterial microbiome may be much higher than our results suggest. However, nearly 70% of the assigned phages were found to target specific *Drosophila* gut bacteria such as *Lactobacillus*, *Acetobacter*, and *Gluconobacter* [49–51]. This indicates that the phage community recovered is at least in part a true representative of the fly gut virome. Nevertheless, some retrieved sequences could represent food or components of the substrate-associated phage communities, as *D. melanogaster* feeds on microbes that grow on decaying substrate.

The majority of the viral sequences identified in this study potentially belonged to novel phage genera, with only a small percentage sharing amino acid identity with previously known phages. Almost 43% of identified sequences clustered into putative viral clusters, with only 5% deriving from a previously established reference genome based on ICTV (International Committee on Taxonomy of Viruses). The accuracy of the classification of the vConTACT2 algorithm [40] has been shown to be high, over 95% based on ICTV, supporting the classification of the sequences in this study. This finding indicates that the *D. melanogaster* microbiome is associated with a diverse range of phages, and that the current knowledge of phage diversity is limited. The fact that we lack full knowledge of phage diversity is further reinforced by similar observations in previous studies on honeybees [47] the human gut [52], and soil phages [53], which also found that only a small percentage of their sequences matched previously identified phages (less than 12%). These findings underscore the importance of continued efforts to study and understand the diversity of phages associated with different host organisms.

Phages are known to facilitate genetic material transfer in host communities, and can affect host phenotypes [54], suggesting that phage-associated proteins may play important roles not just in their life cycles but also in their host. The limited information available to categorize their functions highlights our incomplete understanding of these processes. Nevertheless, the prokaryotic viral-associated proteins that could be assigned to a function were predominantly related to metabolic and DNA processing pathways, in this study. This finding is consistent with observations in honeybees [47]. It indicates that the prokaryotic virome associated with the insect gut includes a vast range of functional genes related to metabolism, which may directly affect the gut ecosystem.

Phages are also shown to contribute to the evolution of their hosts [55] , particularly in the rewiring of metabolic pathways to meet their requirements [56, 57]. Thus, the presence of important metabolic pathways (such as energy, vitamin, carbohydrate, and amino acid metabolisms) in the *Drosophila*-gut-associated virome, similar to observations in honeybees [47] and lizard [3], suggests the phages have the potential to modulate the metabolic state rather than solely rely on the host for resources. On the other hand, some pathways represented in the phages, particularly those related to genetic information processing and nucleotide metabolism, may reflect the phages’ rewiring strategy.

In the context of phage-host interaction, the presence of accessory metabolic genes (AMGs) can determine the success of phage proliferation [5]. This is shown by the prevalence of the dcm gene, which encodes a methyltransferase that protects viral DNA from host restriction enzymes [58, 59]. Similarly, the 7-cyano-7-deazaguanine synthesis genes found also in the *D. melanogaster* bacteriophages offer phage protection against host defenses [60]. Our analysis shows that AMGs in *D. melanogaster* primarily encode products for amino acid, cofactor and vitamins and folding, sorting and degradation. These findings are consistent with previous studies on human and urban environments [2]. We speculate that these AMG functions not only reflect the needs of the phages, but also the environment and the host [53]. Such symbiotic interactions are critical to maintaining the balance of the gut ecosystem and sustaining the health of *D. melanogaster* individuals [61, 62].

Interestingly, our investigation also revealed the presence of pathways for biofilm formation, quorum sensing, and bacterial secretion system pathways within the *D. melanogaster* viral communities. These findings suggest that viruses may interfere with their bacterial hosts’ processes and even influence other bacteria in the same ecosystem [47]. Moreover, the detection of secondary metabolites, terpenoids and polyketides metabolic pathways further indicates an intriguing interaction between phages and hosts. In light of these discoveries, we emphasize that bacteriophages play a critical role in microbial metabolism, and a balanced interaction between them is essential to sustain the continuum of the gut ecosystem. Ultimately, this intricate interplay between phages and bacteria indirectly influences the health and development of *D. melanogaster* individuals.

### DNA viruses of *Drosophila*

DNA viruses of *Drosophila melanogaster* are thought to be relatively rare [17], and consistent with this we found only four viruses that are likely to infect the fly rather than its microbiome. These included Drosophila Kallithea nudivirus, Drosophila Esparto nudivirus, Drosophila Linvill Road densovirus, and Drosophila Vesanto virus.

Drosophila Kallithea nudivirus is a large dsDNA virus that is closely related to a number of other known *Drosophila* pathogens and has been shown to be pathogenic in adult flies [15, 17, 58, 63]. We found that *Drosophila* Kallithea nudivirus had high prevalence across our four sampling sites, with infection rates exceeding 50% of individuals. This is consistent a previous metagenomic study of DNA viruses infecting European *D. melanogaster*, which also reported that *Drosophila* Kallithea nudivirus infects more than half of populations [17].

Drosophila Vesanto virus was the second most common virus of *Drosophila*, being detected in more than 20% of the individuals. Here, using data from single flies, we detected only 10 out of 12 previously described segments of Drosophila Vesanto virus, and we observed distinct variation in the detection of the segments—with segment 11 appearing extremely rare. Based on large pool-sequencing datasets, [17] previously proposed three hypotheses to explain this pattern: that there are 12 segments with extreme variation in copy number; that there is a reassorting community of segments in which some are optional or satellites; or that Vesanto is not a single virus, and multiple viruses appeared in each of the pools presented there—perhaps infecting a component of the cellular microbiome. The very strong association we see between the majority of the segments (and the absence of shared eukaryotic microbiome from some affected individuals) strongly suggests that these are indeed segments of the same virus, as argued by [17] on the basis of a single infected lab line. The complete absence of two segments from all affected flies further confirms that some segments are indeed optional or satellites. However, the lack of negative correlations among most segments, along with the generally consistent pattern of copy-number variation, suggests that in general (apparent) absences are likely driven by the relatively low titre of some segments, rather than segments being optional or homologous being able to substitute for each other. Such variation in genome copy number among infections may hint that Vesanto virus has a multipartite lifestyle [64, 65].

## Data and code availability

Sequencing data will be publicly available at the NCBI upon manuscript publication. The codes for recovering of all the contigs and constructing vMAGs, pangenomic analysis, and visualization are available from the GitHub repository (https://github.com/Minamehr/Virus-Drosophila).

## Acknowledgements

The authors gratefully acknowledge the support of the European Drosophila Population Genomics Consortium (DrosEU), funded by a Special Topics Network (STN) grant from the European Society of Evolutionary Biology (ESEB). Additionally, we appreciate the support from the state of Baden-Württemberg through bwHPC. The authors would like to extend their gratitude to Dr. Mehregan Ebrahimi for his assistance in data analysis.

