## Supplementary Figure for "A diverse gut virome from *Drosophila melanogaster*"

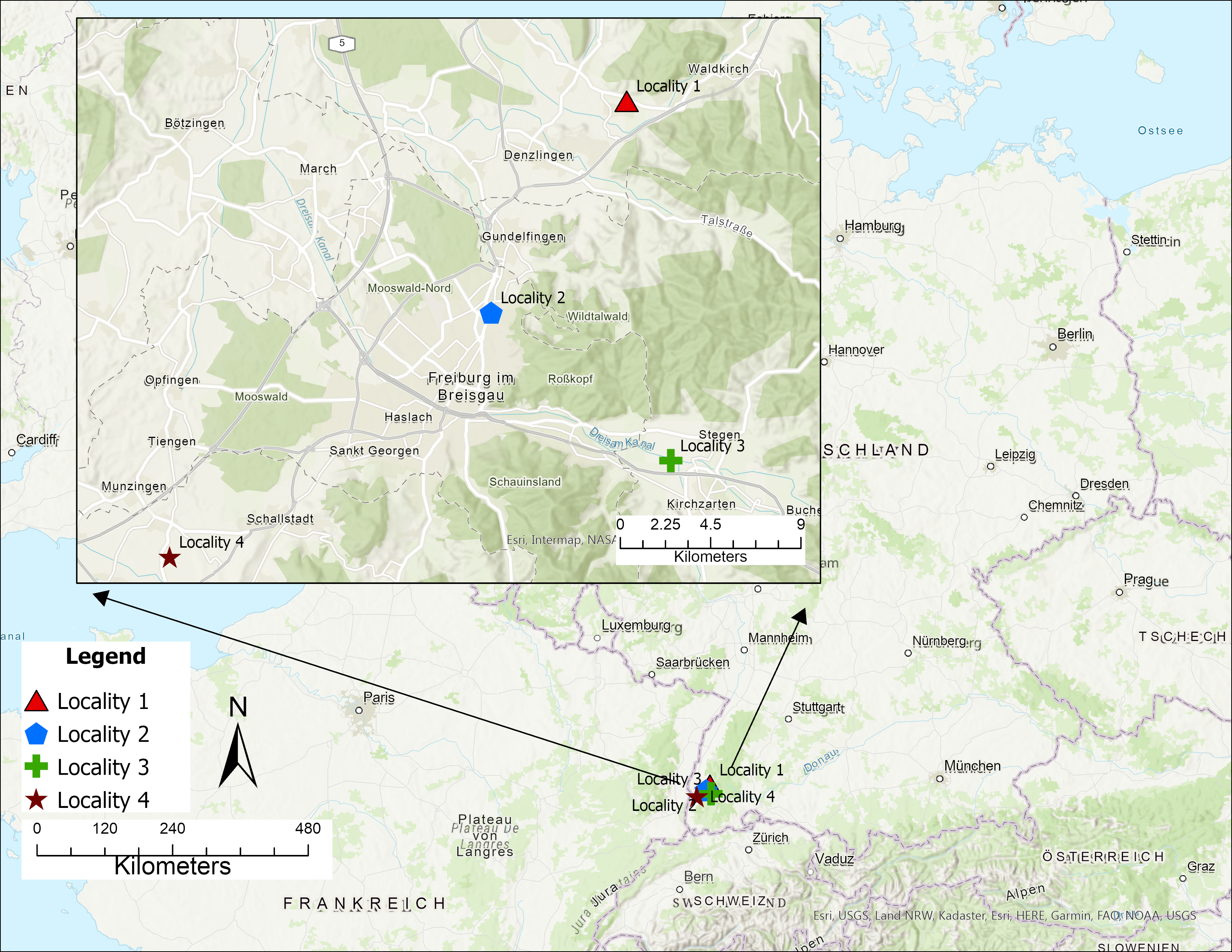

Figure S1. A map representing four sampling locations of *Drosophila* *melanogaster*. We used a mixture of apples and cherry as bait and placed them in multiple 1.5-liter PET bottles at each sampling site for several days. The final data set included 851 metagenomic sequences from 347, 19, 123, and 362 fly samples from sampling location 1, 2, 3 and 4, respectively.

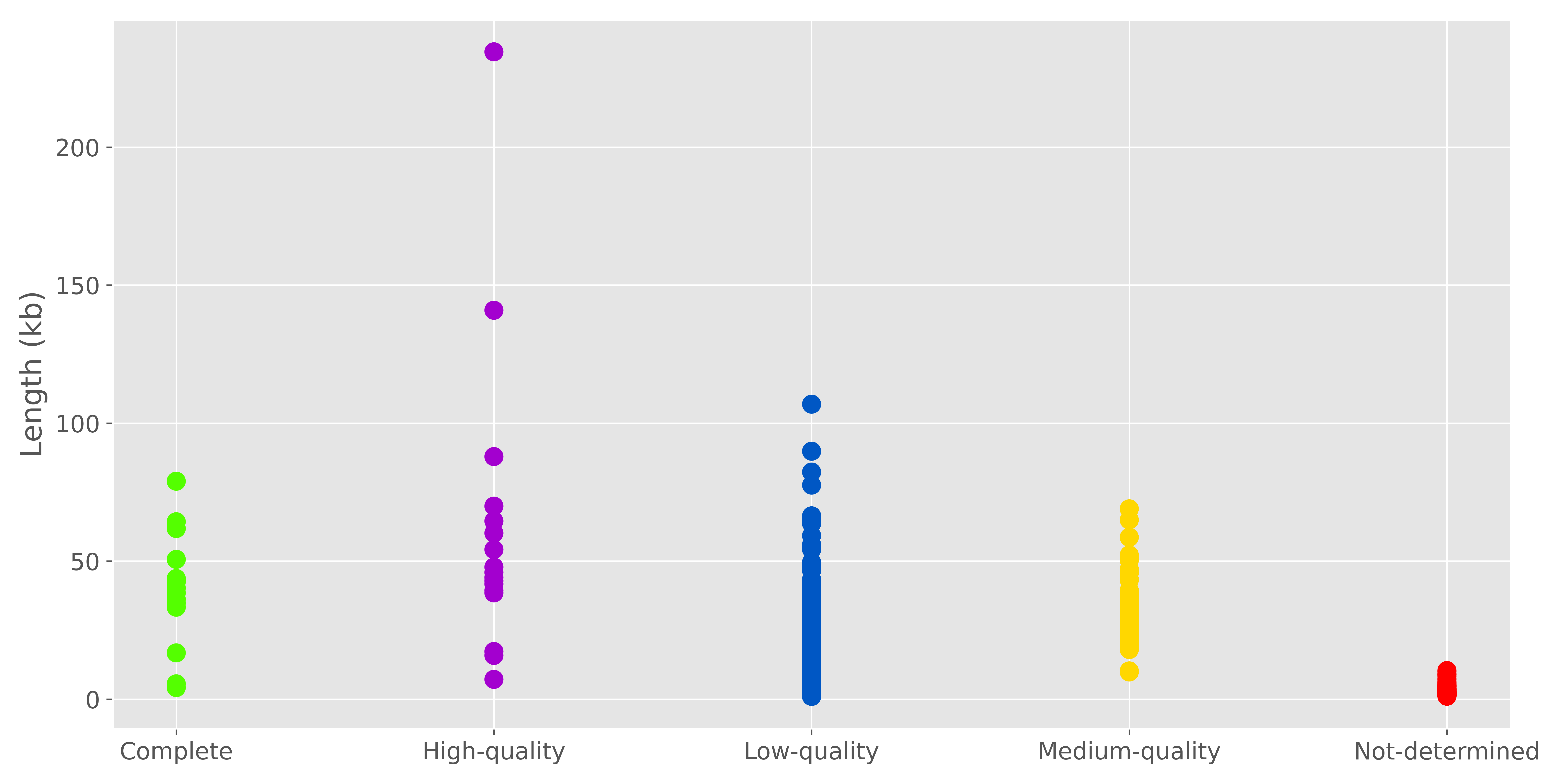

Figure S2. Estimated viral genome qualities as a function of contigs lengths.

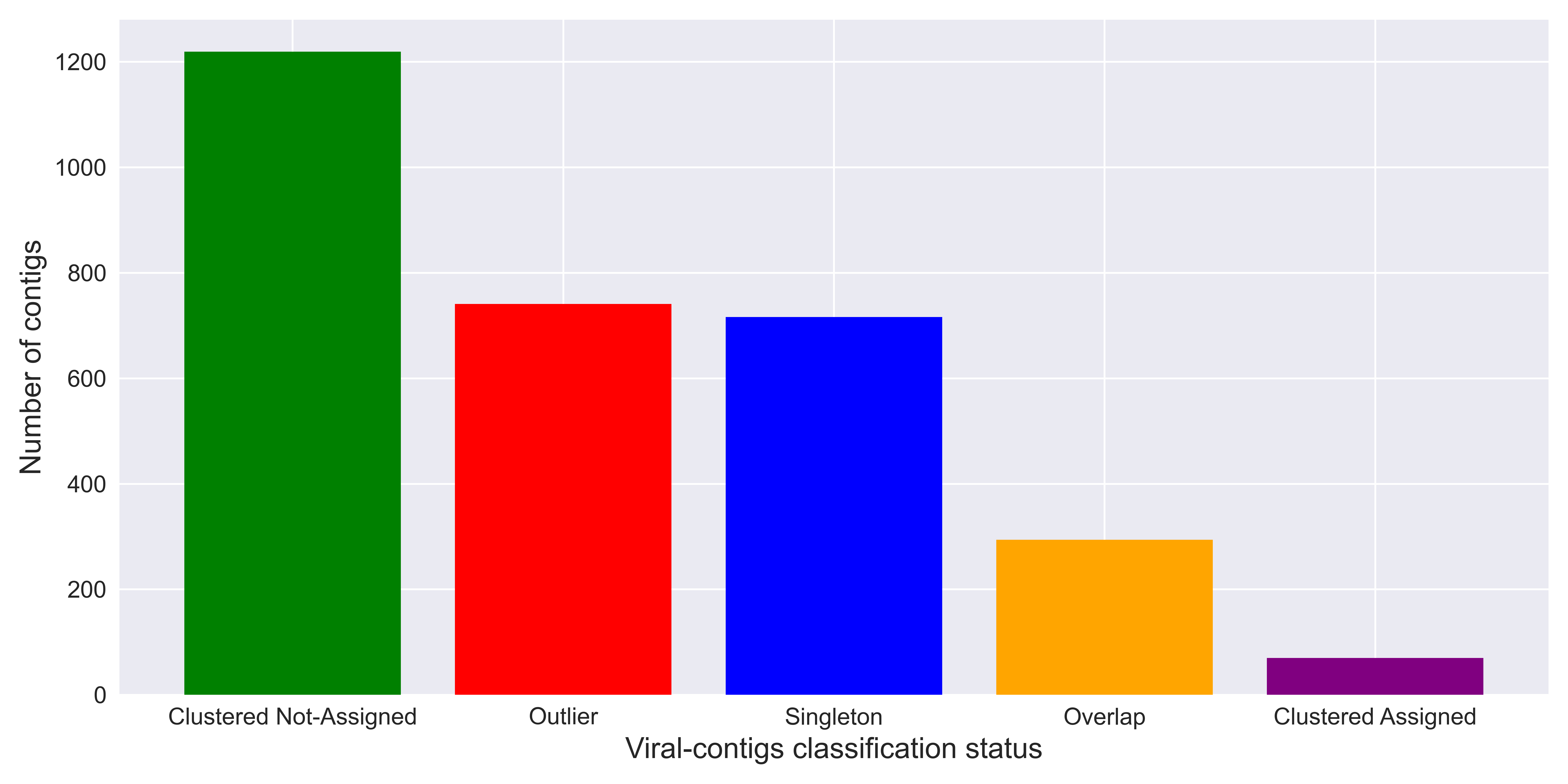

A

B

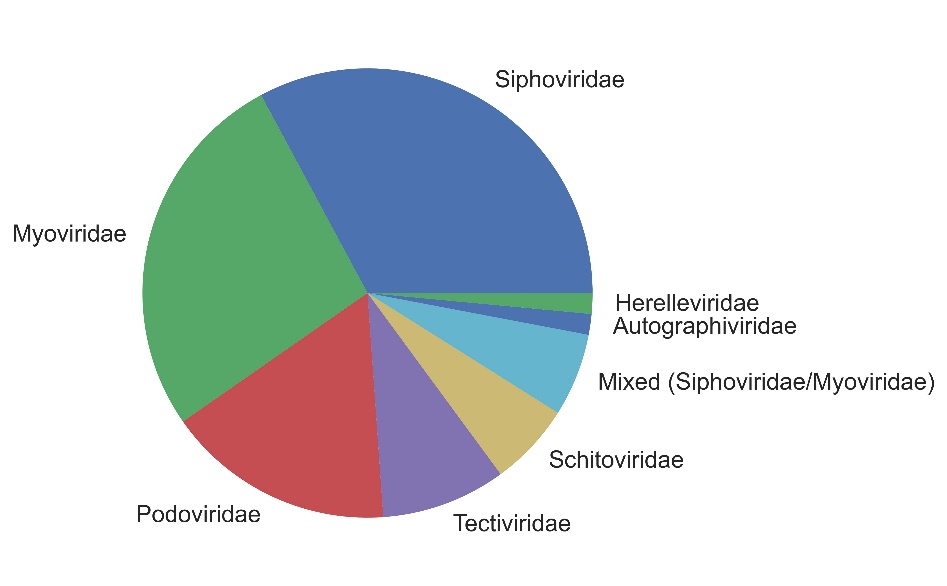

Figure S3. An overview of the recovered viral contigs classification in association with previously described viruses from Refseq database using vConTACT2. A: Reprecenting classification status,

“Clustered Assigned” indicate recovered contigs falling into clusters contained reference sequences; “Clustered Not-Assigned” means recovered contigs clustered without reference sequences. B: Referring to “Assigned Viral contigs” at the family level.

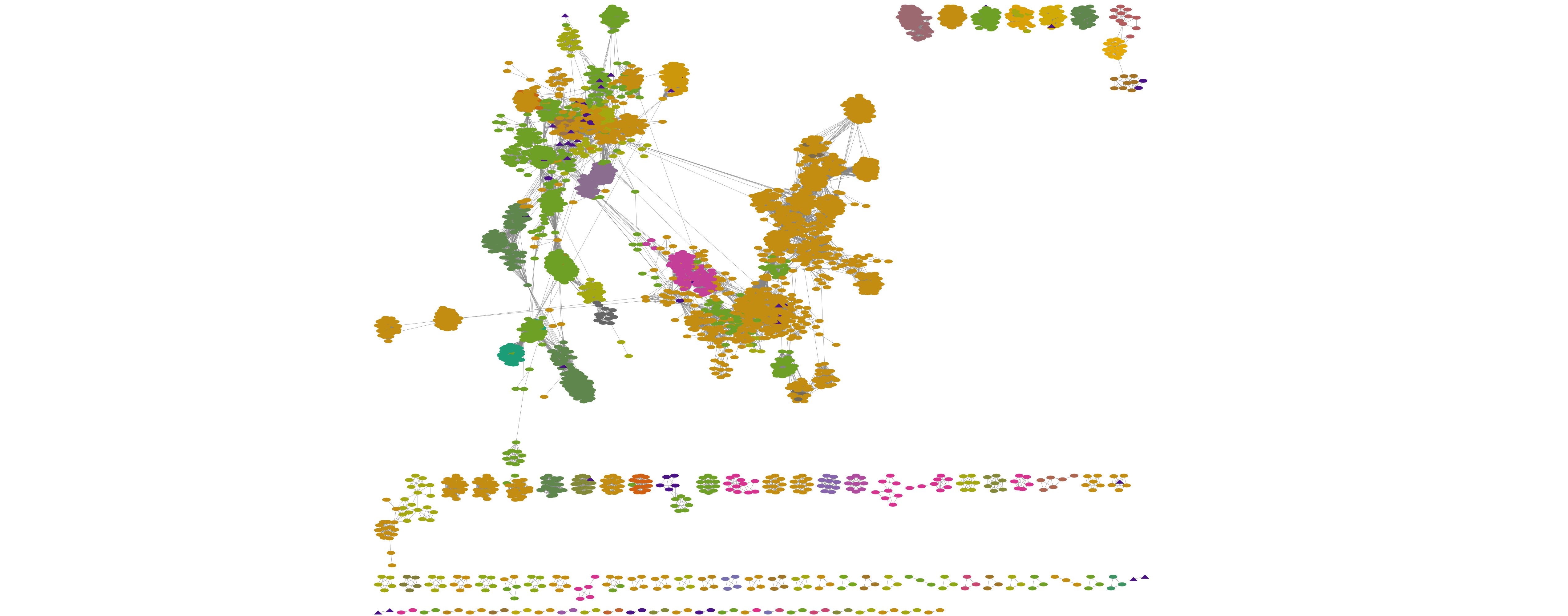

A

B

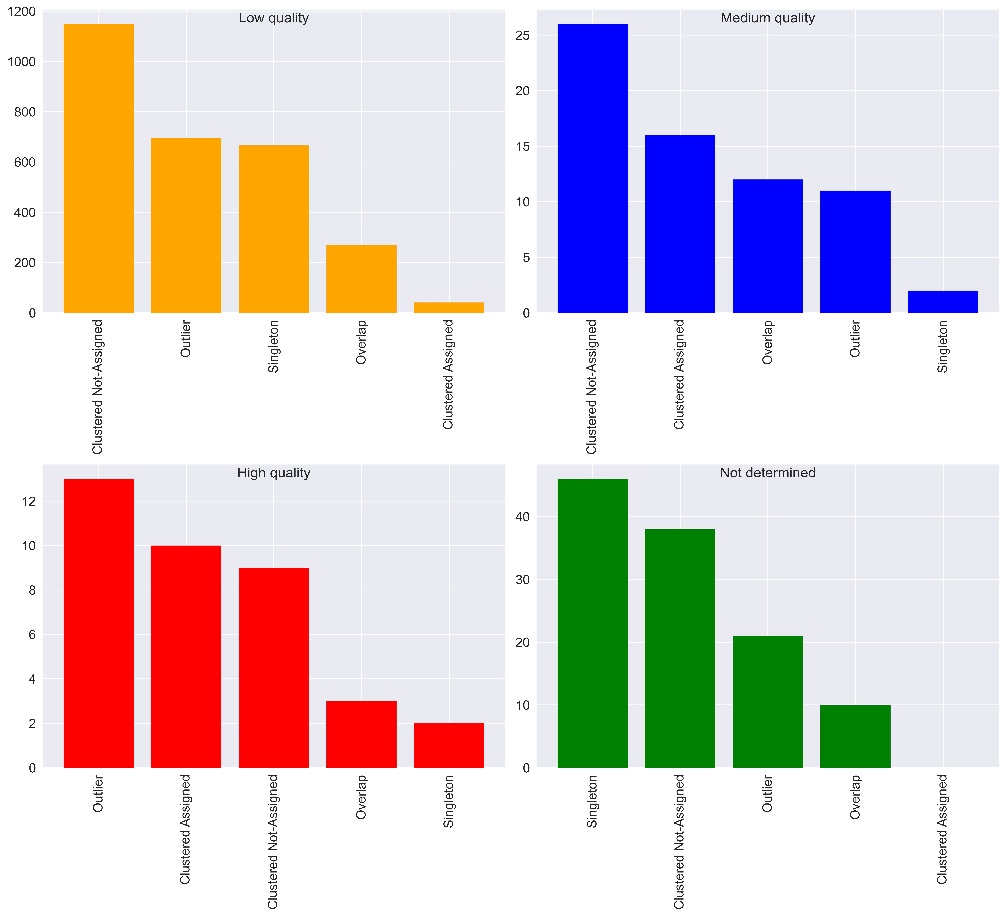

Figure S4. Protein similarity and network analysis constructed with vContact2 and Cytoscape. A: Protein clustering network of the recovered High-quality viral sequences from *D. melanogaster* (purple) compared to previously described viruses from Refseq (other colors). B, C, D and E respectively indicate the classification status of the high, medium and low quality and not-determined quality viral sequences.

C

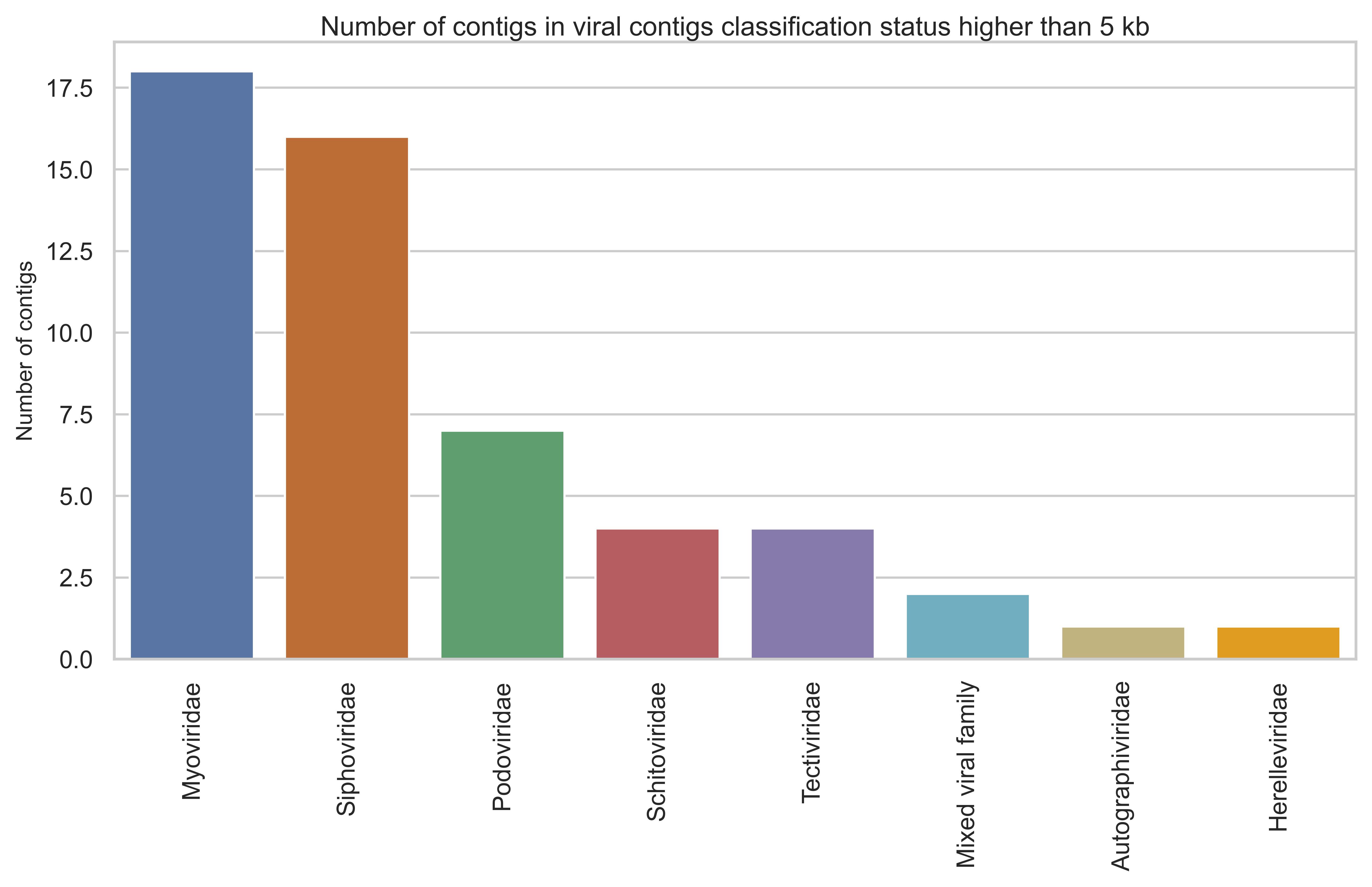

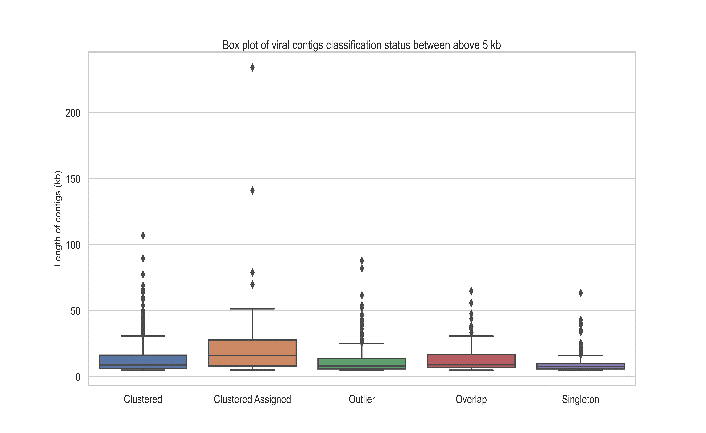

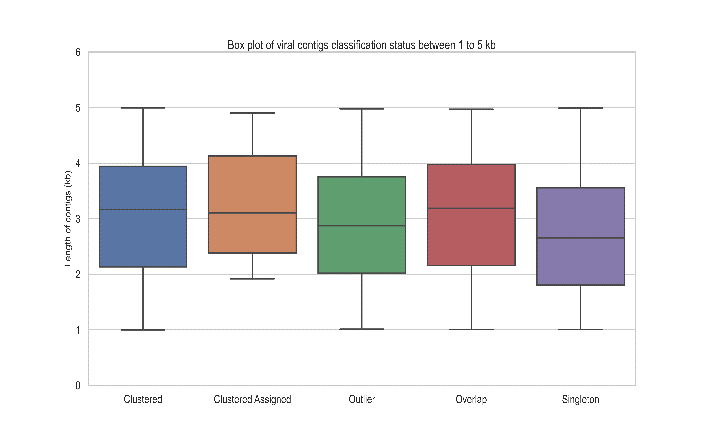

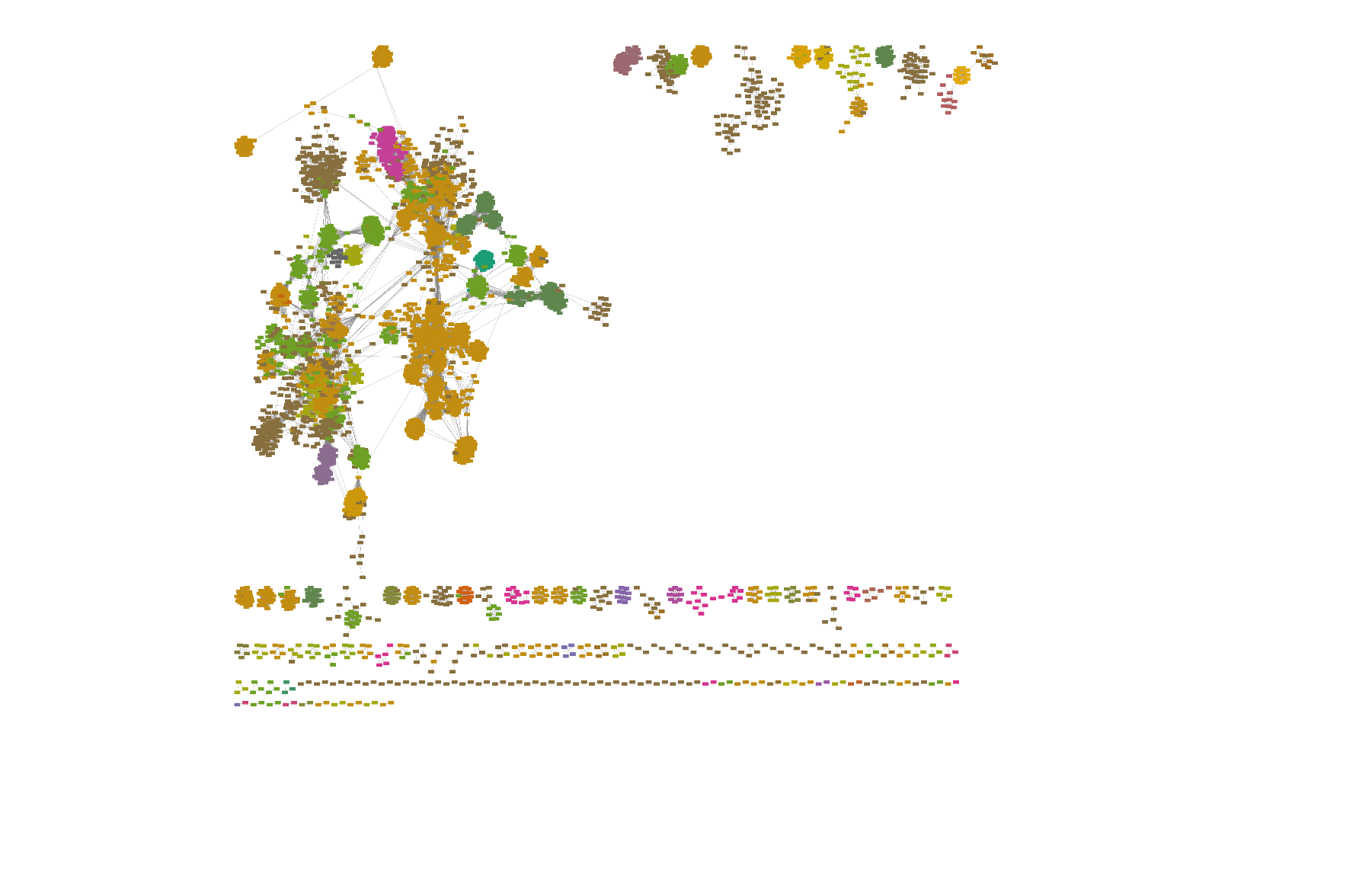

A

B

D

C

Figure S5. An overview of the recovered viral contigs classification status in association with previously described viruses. A: Boxplot represent the classification status of the short recovered viral contigs (<5 kb) using vConTACT2. B: Boxplot represent the classification status of the large recovered viral contigs (>5 kb) using vConTACT2. C: Number of clusters that includes both confidently clustered large contigs and viral reference sequences. D: The network clustering presenting the retrieved clustered putative large (>5 kb) prokaryotic viruses (dark brown) and the viral family of Refseq (other colors).

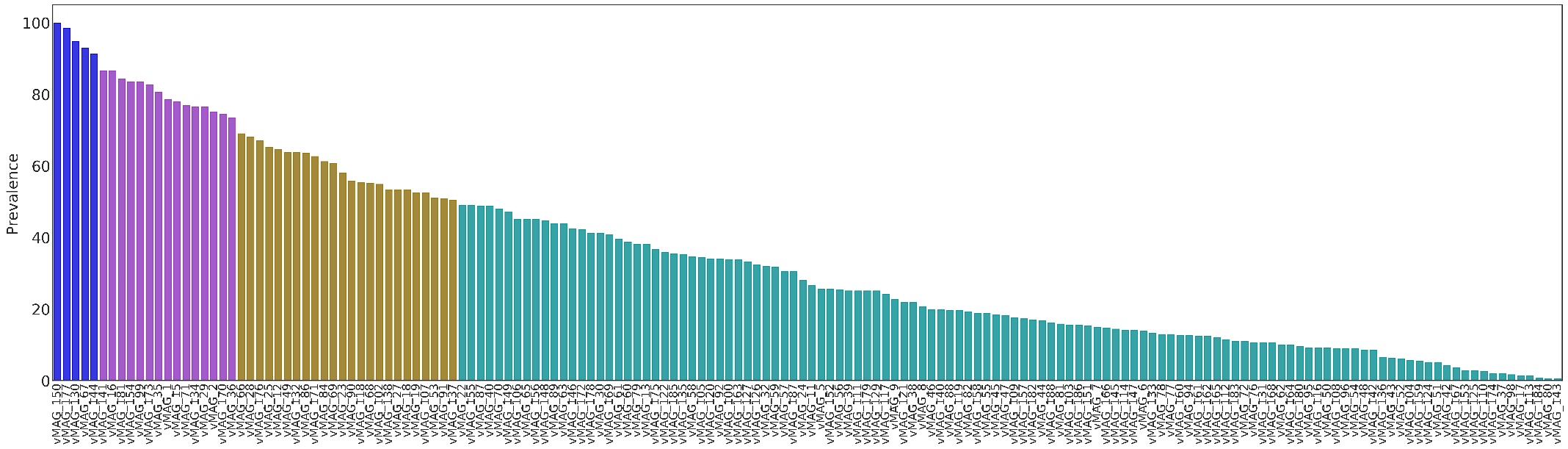

Location 1

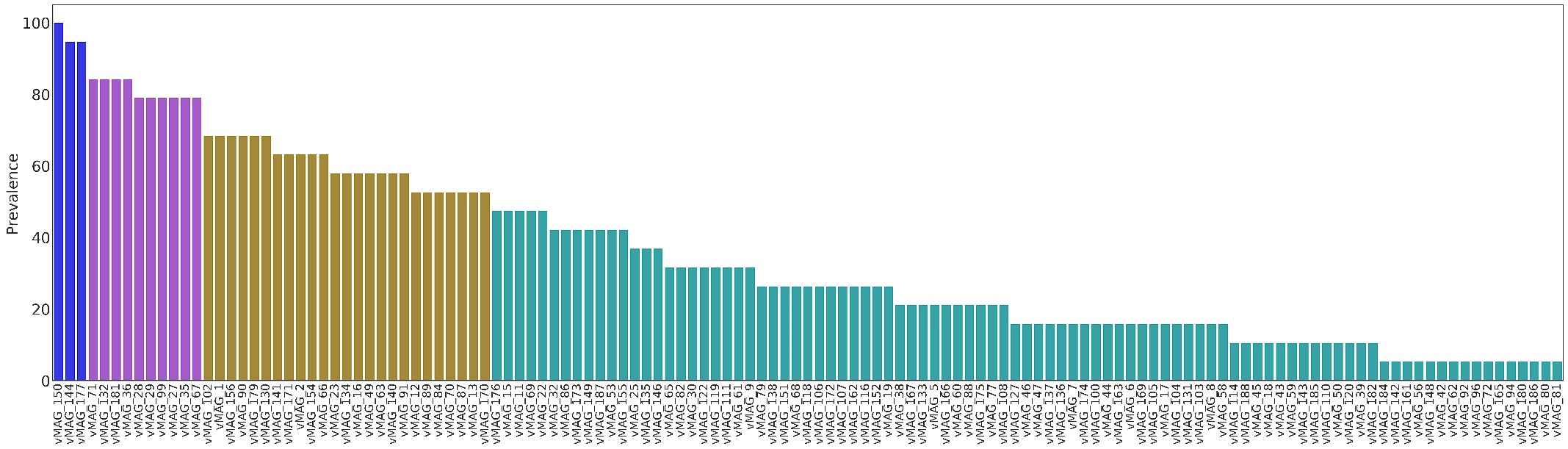

Location 2

Figure S6. The prevalence of each vMAGs (prevalence <50%= green, 50<70%= brown, 70<98%=purple, and 99%< showed with blue color) in each sampling location were shown with the bar charts separetly.

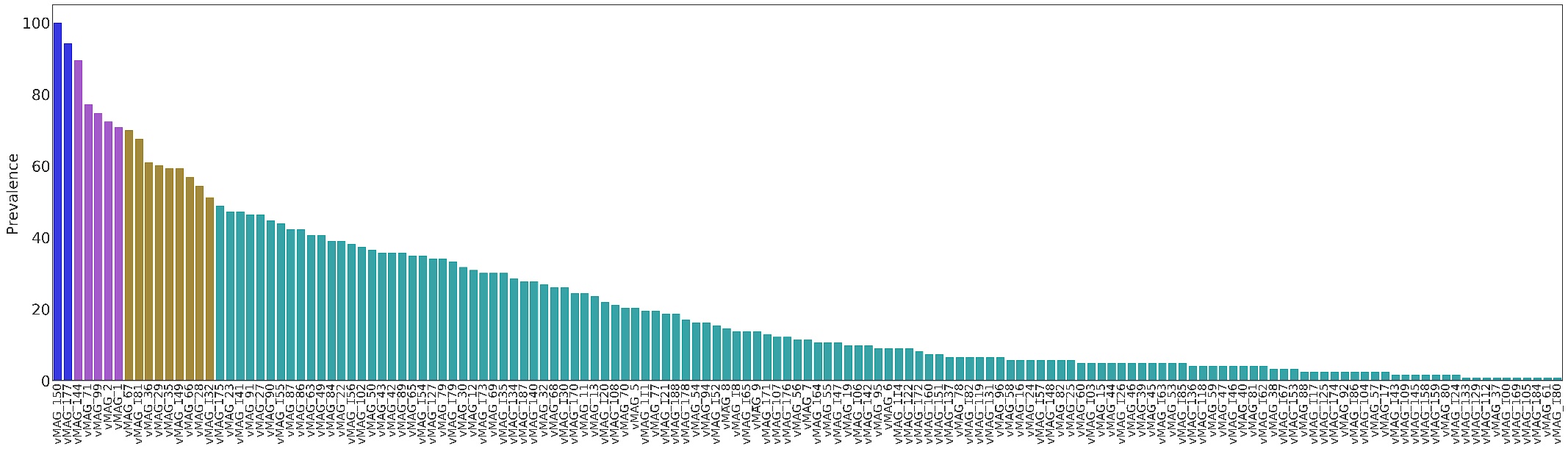

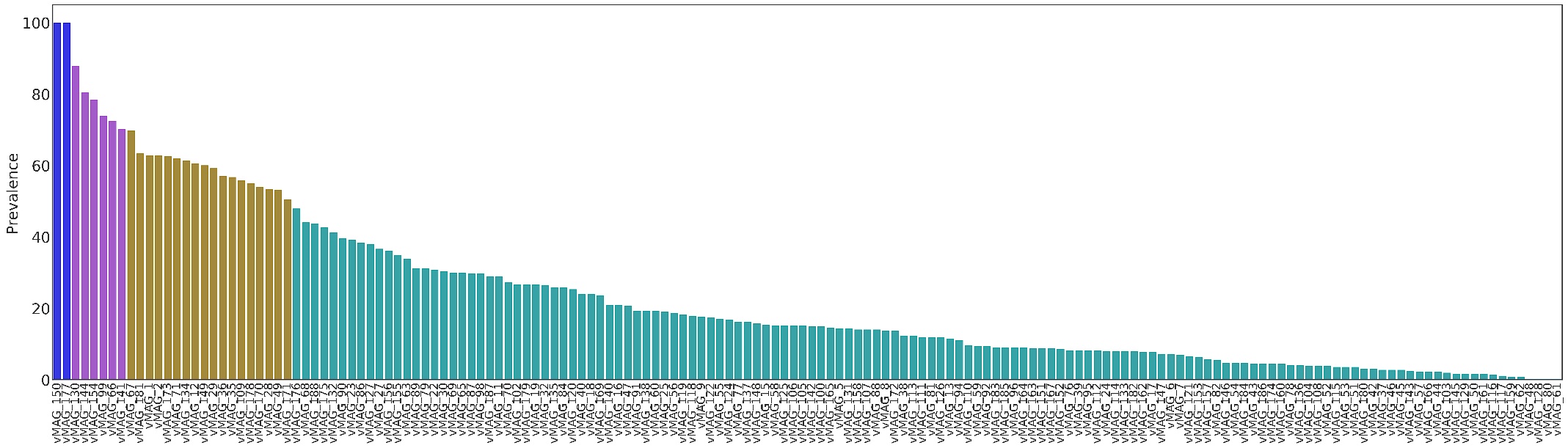

Location 3

Location 4

Continued Figure S6. The prevalence of each vMAGs (prevalence <50%= green, 50<70%= brown, 70<98%=purple, and 99%< showed with blue color) in each sampling location were shown with the bar charts separetly

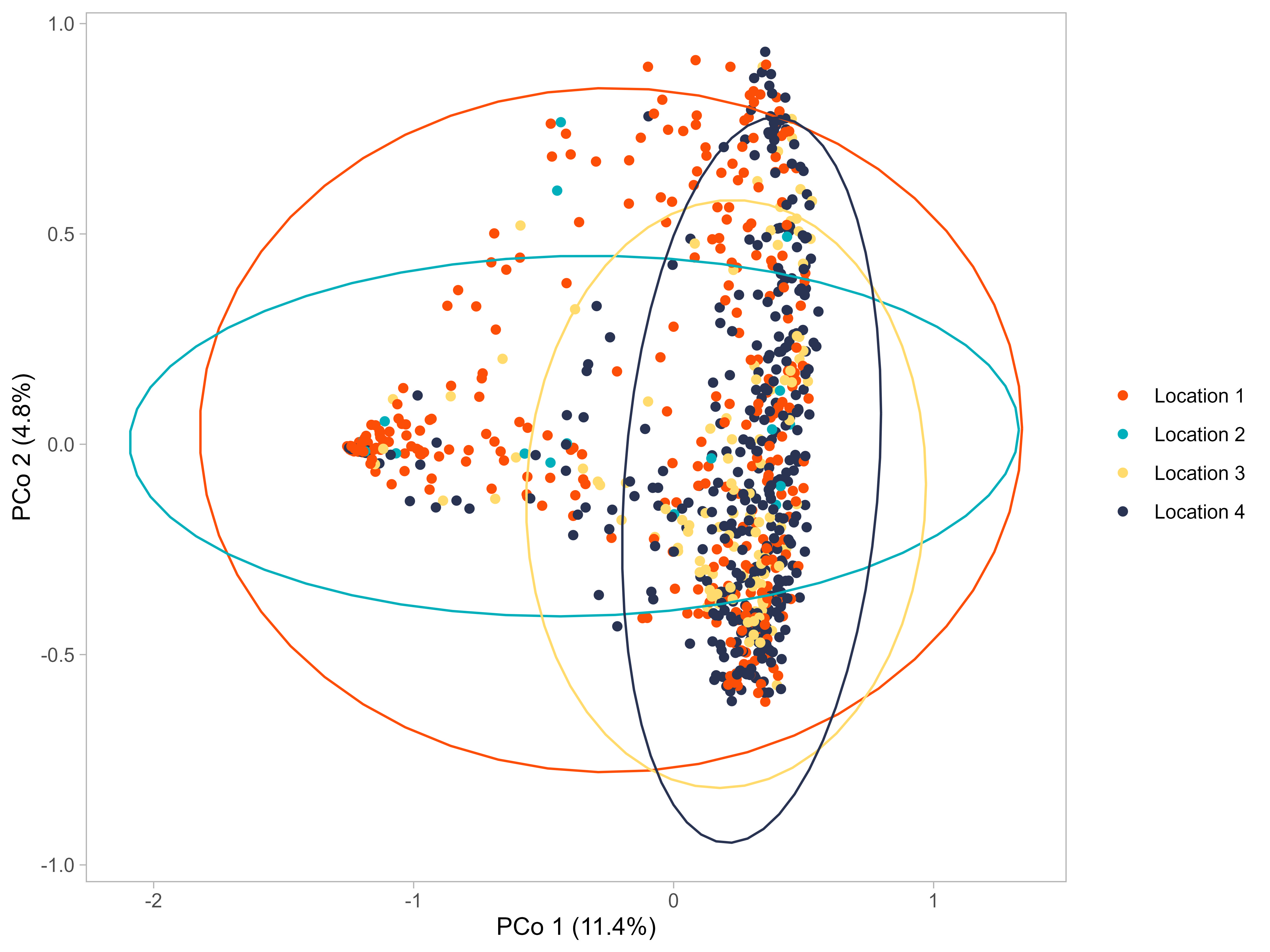

Figure S7. Principal component analysis using Bray–Curtis dissimilarity and based on unbinned viral sequences. P-values of Betadisper was P = 0 .001, also P-values and R squared values from the Adonis2 (with 10,000 permiutation) were P= 9.999e-05 and R= 0.0752, respectively.
